# Genetic Diversity of *Dianthus* Plants Revealed by SRAP Markers

**DOI:** 10.1101/2021.01.04.425199

**Authors:** Jing Wang, Jian Li, Gensheng Shi, Huazheng Hao, Zhongrui Ji, Zhongxiong Lai, Guoming Xing

**Affiliations:** The Industrial Crop Institute, Shanxi Agricultural University, Fenyang 032200, Shanxi, China; Institute of Horticultural Biotechonology, Fujian Agriculture and Forestry University, Fuzhou 350002, Fujian, China; College of Horticulture, Shanxi Agricultural University, Jinzhong 030801, Shanxi, China

**Keywords:** *Dianthus* species, genetic relationship, interspecific, intraspecific, habitats

## Abstract

*Dianthus* is a valuable genetic resource for flower breeding. However, study on the genetic similarity of *Dianthus* plants is rare. In this work, 20 pairs of SRAP markers were used to analyze genetic diversity of 44 *Dianthus* plants, including 13 lines of wild *Dianthus chinensis* L., 7 lines of wild *Dianthus superbus* L. and 24 commercial varieties including *Dianthus caryophyllus* L., *Dianthus plumarius* L. and *Dianthus barbatus* L.. Results showed that precise interspecific and intraspecific genetic diversity was provided in *Dianthus* plants by using 20 pairs of SRAP molecular markers. The interspecific genetic diversity of *Dianthus* plants was much abundant and the intraspecific genetic difference of wild *Dianthus* species was related to their geographical distribution and habitats. In this work, theoretical basis and technical support were provided for crossbreeding and molecular mechanism research of *Dianthus* plants.

## Introduction

*Dianthus* plants are herbs in Caryophyllaceae. About 600 kinds of *Dianthus* species are recorded worldwide, most originated in Europe and Asia, and a few in America and Africa. In China, there are 17 species, 1 subspecies and 9 variants, most of which are grown in the northern grassland and mountainous grassland[1, 2]. *Dianthus* plants all have very significant application value because of their colorful flowers, long flowering period, air purification ability, adaptability to abiotic stress, and so on. *Dianthus caryophyllus* L. is one of the four major cut flowers worldwide and plays an important role in the flower market[3]*. Dianthus plumarius* L. is an important environmental protection and greening plant, because of its strong reproductive capacity and rapid growth speed which can effectively prevent soil erosion and soil erosion [4–6]. *Dianthus chinensis* L. and *Dianthus Superbus* L. are used as herbal medicine for its function of reducing fever and diuresis, and also as insecticidal pesticides[7–9]. *Dianthus barbatus* L. is also an important source of cut flowers and a greening plant. It can purify the air by absorbing harmful gases such as sulfur dioxide and chlorine[10]. Therefore, *Dianthus* is a valuable genetic resource for flower breeding.

It is a common breeding strategy to improve plant characters for *Dianthus* to by aggregating excellent traits from different species through interspecific hybridization[11, 12]. Interspecific genetic difference directly affects efficiency of interspecific hybridization. As a powerful biological research tool, molecular markers are widely used in hybrid breeding, germplasm identification, variety protection and molecular mechanism research[13–16]. Sequence-related amplified polymorphism (SRAP) marker is a popular molecular marker for its simplicity of development, stability of amplification and efficient discrimination[17–19]. It effectively identifies genetic relationship of plants which can provide breeders a better reference for making a crossbreeding plan.

However, study on the genetic similarity of *Dianthus* plants is rare. In this work, SRAP markers were used to analyze genetic diversity of 44 *Dianthus* plants in order to explore the genetic relationship among *Dianthus* species and provide theoretical basis and technical support for crossbreeding and molecular mechanism research of *Dianthus* plants.

## Materials and Methods

### Plant Materials

44 *Dianthus* plants were used in this work including 13 lines of wild *Dianthus chinensis* L., 7 lines of wild *Dianthus superbus* L. and 24 commercial varieties including *Dianthus caryophyllus* L., *Dianthus plumarius* L. and *Dianthus barbatus* L. (Table S1). Collection area information of wild species was given in Table S2.

### DNA Extraction

Young-leaf tissue from samples was ground in liquid nitrogen to a fine powder and total genomic DNA was extracted using the modified CTAB method. DNA quality testing was performed by 1% agarose gel electrophoresis.

### SRAP Analysis

88 primer pairs were used in this work which consisted of 8 forward primers and 11 reverse primers. The SRAP reaction mixture (total volume = 20 μL) contained 40 ng DNA, 2 μL 10 × buffer, 1.5 mM MgCl_2_, 0.5 μL primers, 2 μL dNTPs (2.5 mM), 1.5 U *Taq* DNA polymerase (Fermentas USA). The first step was an initial preheating (5 min at 94 °C). The subsequent 5 cycles each was carried out that consisted of denaturation at 94 °C for 30 s, annealing at 30 °C for 1 min and extension at 72 °C for 80 s. The third step was extension at 72 °C for 10 min, and the next 37 cycles each consisted of the steps: 94 °C for 1 min, 50 °C for 1 min, and 72 °C for 80 s, and a final extension step of 72 °C for 12 min was performed. All PCR reactions were performed in a PTC-100™ programmable Thermal Controller (Bio-Rad, USA). Amplified products were detected by 6% polyacrylamide gel electrophoresis run in 1 × TBE buffer at 120V and photographed after silver staining. Primers were synthesized by Sangon Biotech (Shanghai) Co., Ltd. and primer sequences were listed in Table 1.

**Table 1.**
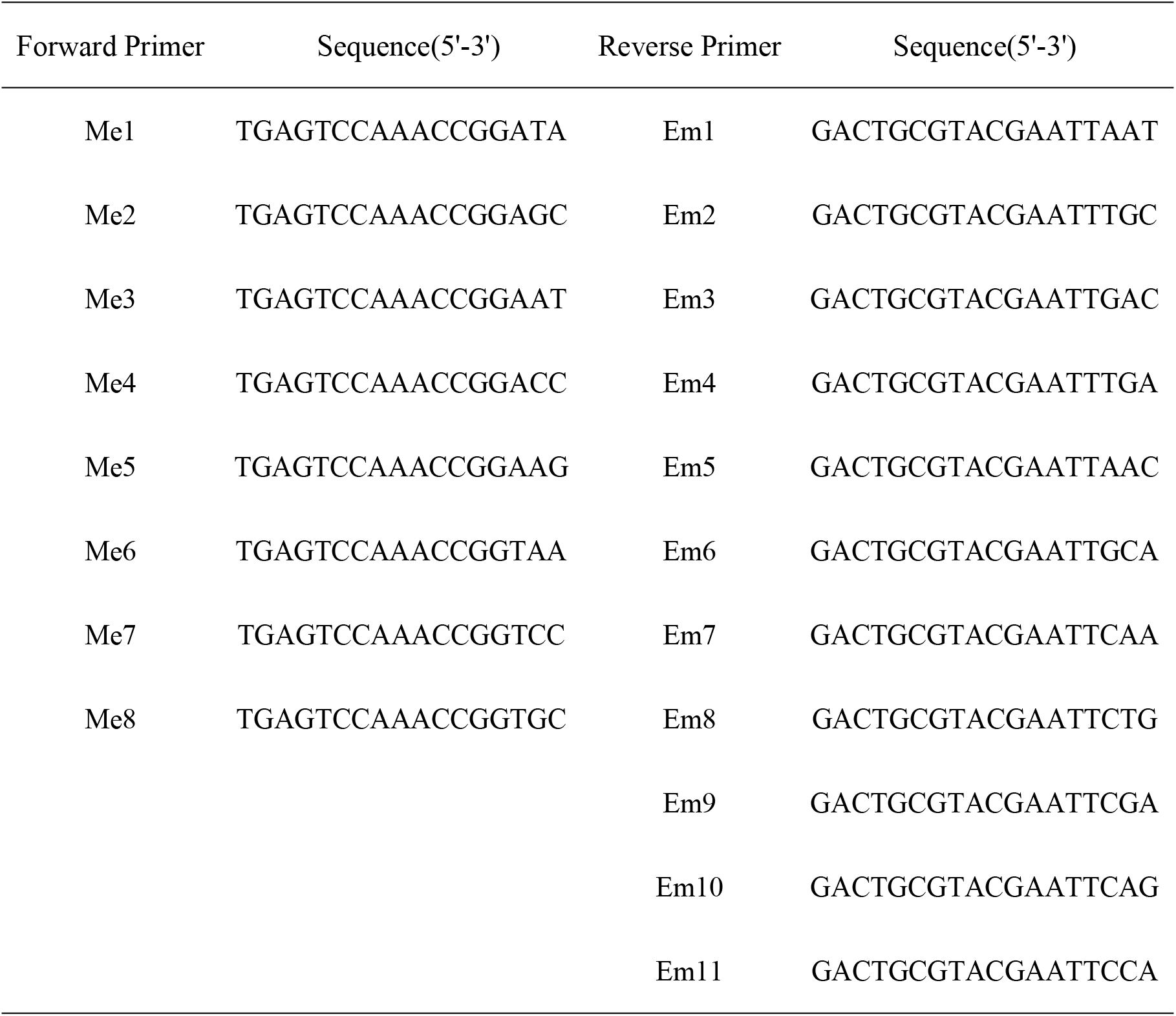
Information of SRAP primers.

### Data Analysis

In the SRAP marker analysis, the banding patterns were recorded as present (1) or absent (0) for each primer pair. The number of bands, number of polymorphic bands and number of distinguishable kinds of each primer pair were counted. The distance matrix was subjected to cluster analysis by the unweighted pair-group method (UPGMA) and the dendrogram was performed by NTSYSpc-2.0 software.

## Results

### DNA Extraction and Testing of PCR Products

Genomic DNA was showed clear bands and undegraded extracted by modified CTAB method which mean the DNA was qualified for genotyping (Fig 1a). Only 20 primer pairs in 88 pairs were amplified successfully (Table 2, Fig 1b).

**Table 2.**
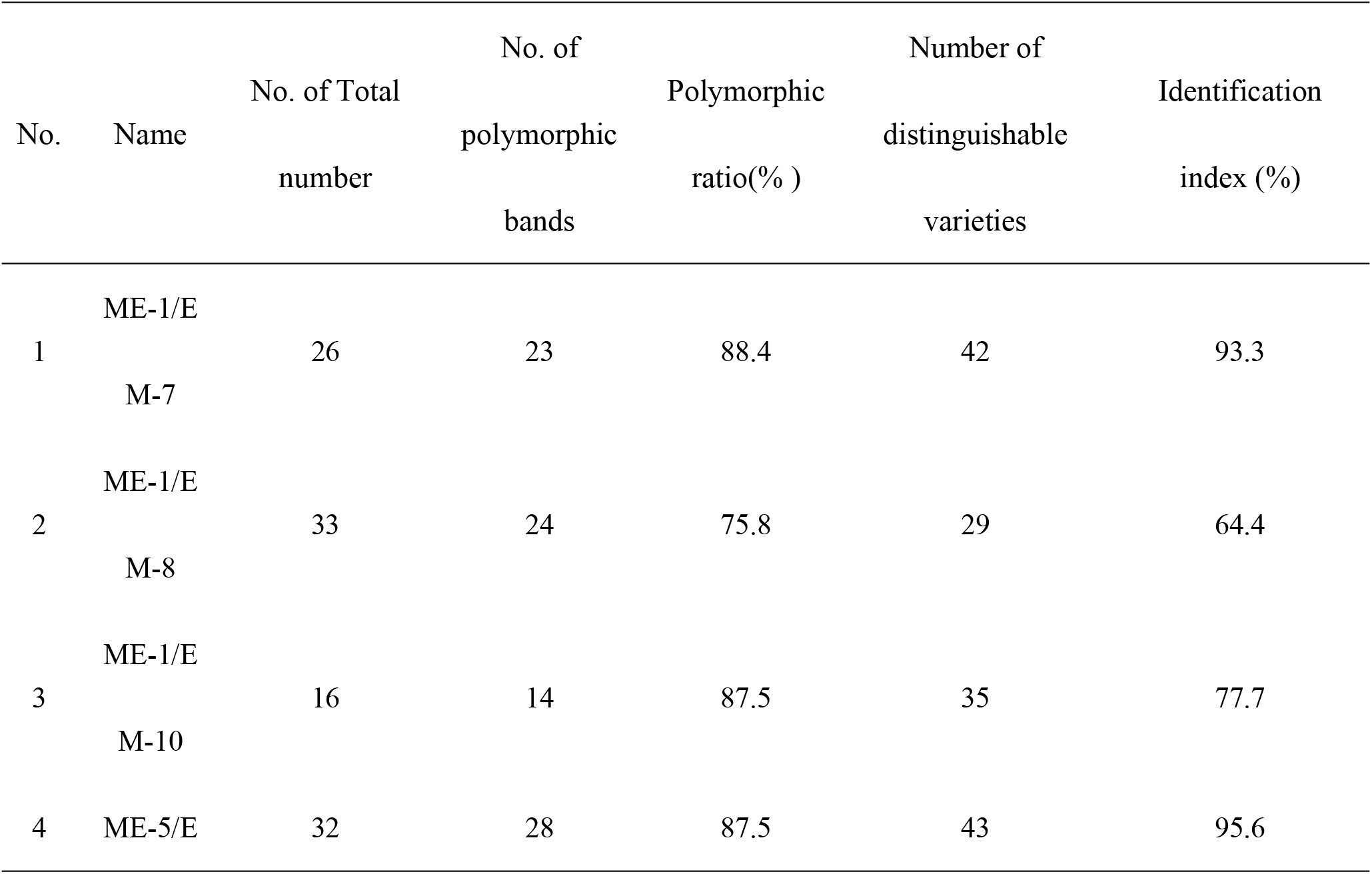
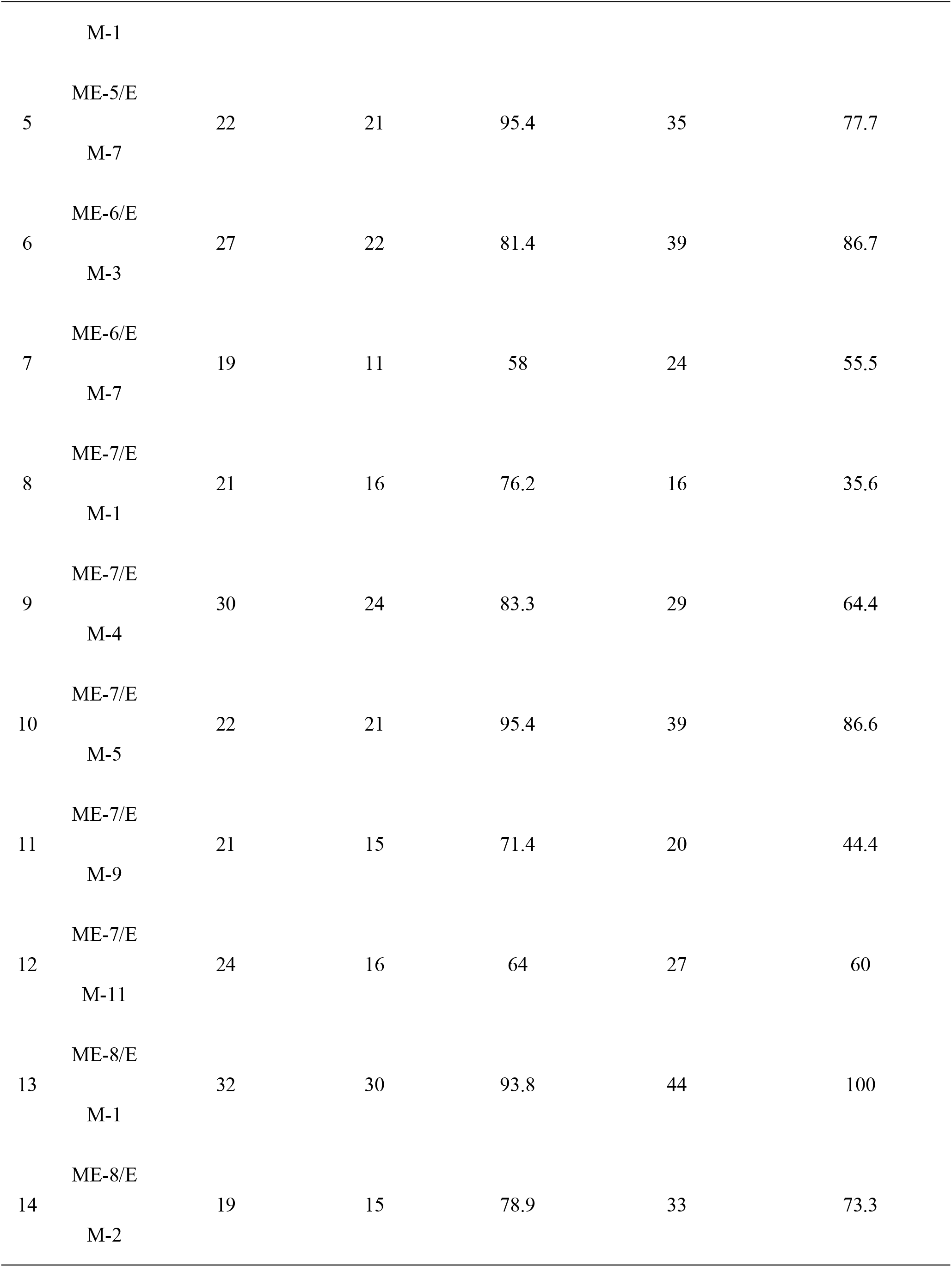
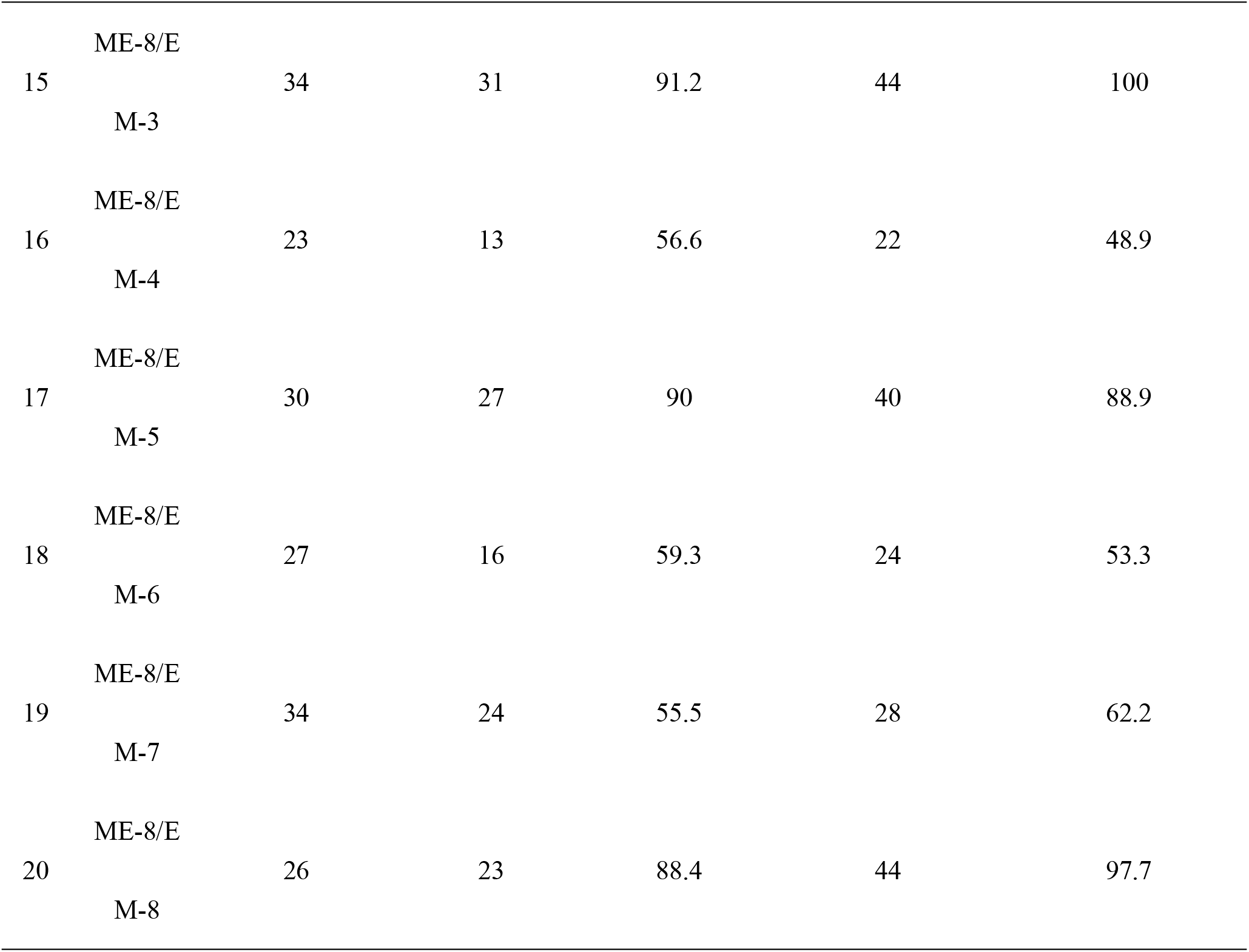
Polymorphism statistics of PCR products amplified by 20 pairs of SRAP primers.

**Fig 1.**
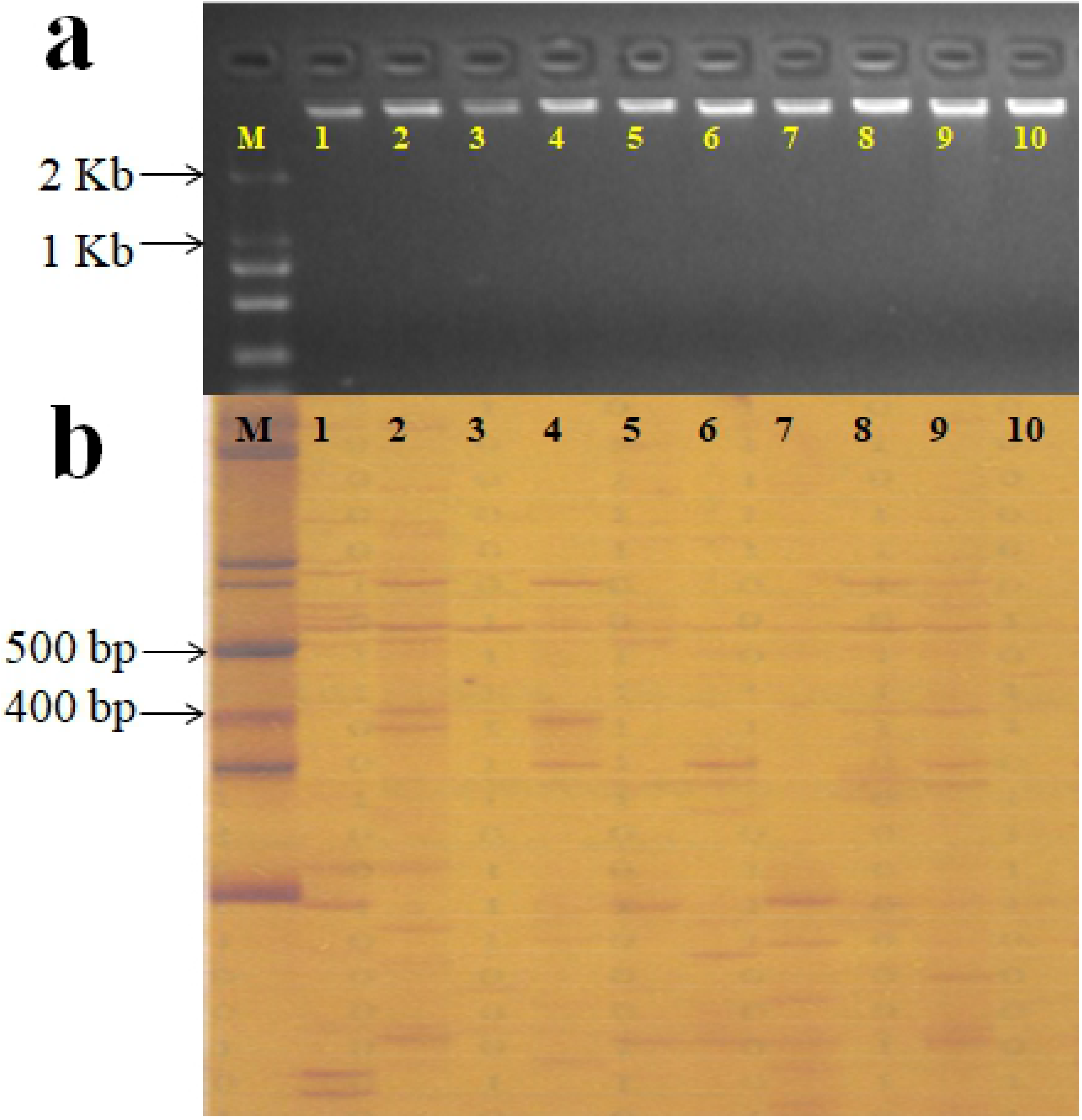
Quality control of DNA and the gel electrophoresis of PCR products. a) M: marker; 1-10 represent agarose gel electrophoresis resulted from 10 DNA samples; b) M: marker; 1-10 represent polyacrylamide gel electrophoresis resulted from 10 PCR products amplified by primer pair Me-8/Em-3.

### SRAP Marker Analysis and Construction of Fingerprint for Samples

In this work, 518 bands were amplified by 20 primer pairs, and the average number of total bands was 25.9. The average number of polymorphic bands was 20.7 and the ratio of it was 78.9%. The identification ability of 20 primer pairs was ranged from 35.6% to 100% (Table 2). Primer pair Me-8/Em-3 showed the highest discrimination, which could distinguish all samples. Fingerprint of each sample amplified by 20 primer pairs was constructed, and the fingerprint of samples amplified by Me-8/Em-3 was showed as following (Table S3).

### Analysis of Genetic Diversity

In this work, 414 polymorphic bands were used for genetic similarity analysis. The Jaccardp’s similarity coefficient ranged from 0.73 to 0.98. 44 samples were divided into 5 groups which were in accordance with the species (Fig 2). In these 5 groups, *Dianthus plumarius* L. showed the minimum genetic similarity with other four species, and the second and the third were *Dianthus caryophyllus* L. and *Dianthus barbatus* L. respectively (Fig 2).

**Fig 2.**
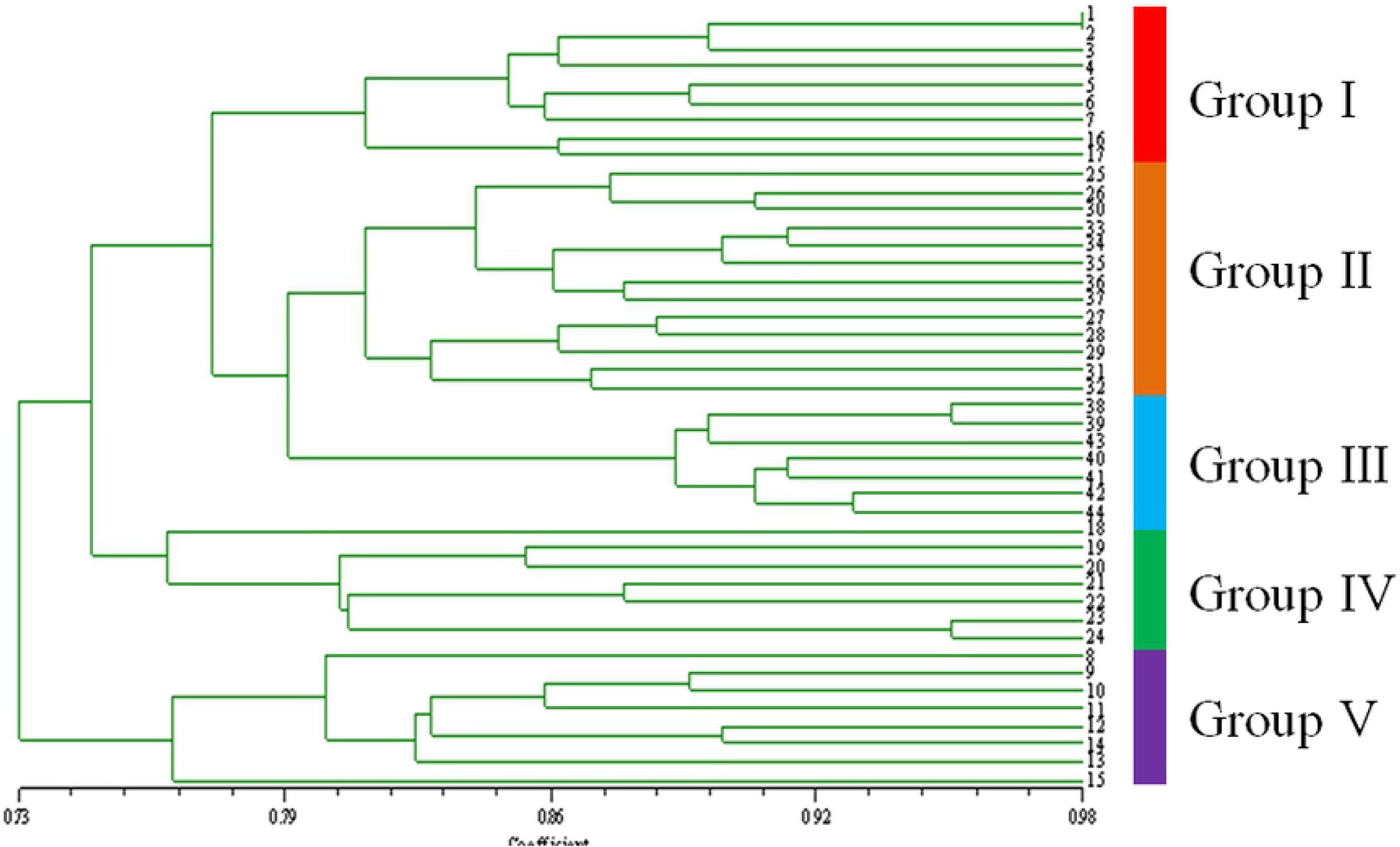
Clustering result of 44 samples by 20 pairs of SRAP markers.

## Discussion and Conclusions

Molecular markers are becoming a powerful tool for plant variety identification and protection, variety selection and molecular mechanism research because of its high detection efficiency and precision. In this work, 20 primer pairs of SRAP markers were screened and showed a high resolution and precise identification in 44 *Dianthus* samples. Genetic distance between these 5 groups showed a significant difference. The genetic similarity of *Dianthus chinensis* L. and *Dianthus superbus* L. was the highest following by *Dianthus barbatus* L., and *Dianthus plumarius* L. showed the lowest genetic similarity with the other four. Phenotypically, *Dianthus plumarius* L. showed the dense clump stem and the other four were the sparse clump stem. Except *Dianthus plumarius* L., *Dianthus caryophyllus* L. showed a higher plant height than the other three.

In wild species, the intraspecific genetic similarity was strongly associated with the habitat. The wild *Dianthus chinensis* L. group was divided into 3 subgroups. The first subgroup was distributed in the south-central Shanxi Province including 25/26/30. The second and the third were distributed in the north Shanxi Province including 33/34/35/36/37 and the central Shanxi Province including 27/28/29/31/32 respectively (Fig 2). The first and the second subgroup located in low latitude showed a higher genetic similarity than the third one located in high latitude (Fig 2, Table S1).Similarly, the wild *Dianthus superbus* L. group was also divided into 2 subgroups, one in the south Shanxi Province including 38/39/43 and another in north-central Shanxi Province including 40/41/42/44 (Fig 2, Table S1). In this work, the intraspecific genetic similarity of wild species was closely related to their geographical distribution and habitats. And the intraspecific genetic similarity of cultivars showed a high relationship with their phenotypic characteristics. For example, *Dianthus barbatus* L. group was divided into 2 subgroups, one including 1/2/3/4/5/6/7 and another including 16/17. The forward could also be divided into 2 groups, pink flower group including and white/red flower group. Furthermore, the pink flower group could also be divided into 2 groups, light pink and dark pink.

In conclusion, the interspecific genetic diversity of *Dianthus* plants was much abundant and the intraspecific genetic difference of wild *Dianthus* species was related to their geographical distribution and habitats. Precise interspecific and intraspecific genetic diversity was provided in *Dianthus* plants by using SRAP molecular markers. In this work, theoretical basis and technical support were provided for crossbreeding and molecular mechanism research of *Dianthus* plants.

## Acknowledgements

This study was supported by TDPAF MFAFF/iPET through Grape Research Project Group.

## Supporting information

**S1 Table. Information of sample species and source**

**S2 Table. Collection area information of wild species**

**S3 Table. Fingerprints of 44 samples established by ME-8/EM-3.**

